# Proteomic changes orchestrate metabolic acclimation of a unicellular diazotrophic cyanobacterium during light-dark cycle and nitrogen fixation states

**DOI:** 10.1101/2024.07.30.605809

**Authors:** Punyatoya Panda, Swagarika J. Giri, Louis A. Sherman, Daisuke Kihara, Uma K. Aryal

## Abstract

Cyanobacteria have developed an impressive array of proteins and pathways, each tailored for specific metabolic attributes, to execute photosynthesis and biological nitrogen (N_2_)-fixation. An understanding of these biologically incompatible processes provides important insights into how they can be optimized for renewable energy. To expand upon our current knowledge, we performed label-free quantitative proteomic analysis of the unicellular diazotrophic cyanobacterium *Crocosphaera subtropica* ATCC 51142 grown with and without nitrate under 12-hour light-dark cycles. Results showed significant shift in metabolic activities including photosynthesis, respiration, biological nitrogen fixation (BNF), and proteostasis to different growth conditions. We identified 14 nitrogenase enzymes which were among the most highly expressed proteins in the dark under nitrogen-fixing conditions, emphasizing their importance in BNF. Nitrogenase enzymes were not expressed under non nitrogen fixing conditions, suggesting a regulatory mechanism based on nitrogen availability. The synthesis of key respiratory enzymes and uptake hydrogenase (HupSL) synchronized with the synthesis of nitrogenase indicating a coordinated regulation of processes involved in energy production and BNF. Data suggests alternative pathways that cells utilize, such as oxidative pentose phosphate (OPP) and 2-oxoglutarate (2-OG) pathways, to produce ATP and support bioenergetic BNF. Data also indicates the important role of uptake hydrogenase for the removal of O_2_ to support BNF. Overall, this study expands upon our knowledge regarding molecular responses of *Crocosphaera* 51142 to nitrogen and light-dark phases, shedding light on potential applications and optimization for renewable energy.

## Introduction

Cyanobacteria are diverse photosynthetic organisms that play a vital role in harvesting solar energy (1). Their ability to produce oxygen contributed to the oxygenation of the Earth’s atmosphere that not only made the Earth hospitable for other life forms that rely on oxygen but also allowed for more complex organisms to evolve (2) (3). The ability to consume CO_2_, harvest solar energy, and convert it into chemical energy allowed them to succeed in an otherwise hostile environment. Consequently, cyanobacteria provide a powerful solution for carbon-neutral energy production, carbon sequestration, and a wealth of targets for metabolic engineering of energy-rich biomolecules (4). Among many unicellular diazotrophs, *Crocosphaera subtropica* (formerly known as *Cyanothece* sp. ATCC 51142) have evolved diurnal rhythms and biological nitrogen fixation (BNF), a process sensitive to oxygen (5–7), making it interesting and metabolically versatile cyanobacteria yet studied. It can maintain an anoxic cytoplasmic environment to prevent oxygen toxicity of the nitrogenase enzymes which are responsible for BNF. The genome indicates a wealth of metabolic potential, in addition to highly active photosynthesis and CO_2_-uptake mechanisms (8). Importantly, it can perform BNF either in the dark or during a prolonged continuous light phase after entrainment of light-dark growth (9, 10) and has evolved as an attractive model to study processes such as photosynthesis, BNF, and carbon sequestration as well as for understanding their circadian and diurnal rhythms (11, 12).

The presence or absence of light and nutrition (carbon and nitrogen), trigger a cascade of gene expression, protein translation, signaling, and shifts in membrane organization that lead to the functional changeover from photosynthesis to BNF, and *vice-versa*. In tandem with these metabolically exclusive processes are a host of other time-dependent processes, which the cell switches based on its internal mechanisms as if anticipating the light-dark cycle, rather than reacting to it (9, 13). Under nitrogen-fixing condition, the cells become filled with large granules between the photosynthetic membranes (9, 10). These granules, known as glycogen granules, contain semi-amylopectin, essential to generate energy through their degradation during BNF (14). The composition and changes of enzyme activity between light-dark transitions for these glycogen granules are still outstanding. An understanding of the mechanistic processes that link between sensing light-dark transitions, protein translation, membrane organization, and signaling provides key insights into how *Crocosphaera* 51142 might be optimized for enhanced renewable energy production. The expression of nitrogenase is limited to nitrate-depleted growth, and in the absence of an external carbon source, nitrogenase expression is strictly restricted to the dark cycle (15–17). However, when cells are grown in the presence of sufficient carbon, such as glycerol, nitrogenase expression occurs during both the light and dark cycles, along with other enzymes related to respiration, glycogen metabolism, and glycolytic/pentose phosphate pathways (15). These observations suggest that enhanced respiration during the light cycle, attributed to excess carbon sources, may create an anoxic cellular condition, and this condition, in turn, facilitates the expression of nitrogenase.

Nitrogenase enzymes are rapidly inactivated or degraded upon exposure to O_2_ (18, 19). *Crocosphaera* 51142 metabolic machinery is cycling daily between O_2_ production in photosynthesis during the day and oxygen scavenging during respiration and BNF at night. Under subjective light, cells also show distinct circadian rhythms of photosynthesis and BNF with peaks every 24 hours. Interestingly, under subjective light, the capacity for photosynthesis is reduced during the period of BNF (20), even though light energy is available. It is possible that the circadian rhythms and transitions between photosynthesis and BNF are correlated not only with intense respiration and nitrogenase activity but also with redox state of the cell through the action of thioredoxin (21) and activation or inhibition of PSI and PSII complexes (20). While previous transcriptomic and proteomic studies have shed light on certain aspects of the regulation of this desperate metabolic process, there is still much to be unmasked about the intricate interplay between nitrate, light-dark cycles, and metabolic processes in cyanobacteria. Proteomic technologies are also constantly improving for greater sensitivity, resolution, and accuracy, allowing in-depth analysis. Taking advantage of such new concepts and improved technologies, we performed quantitative mass spectrometry analysis of *Crocosphaera subtropica* ATCC51142 to determine mechanisms underlying shift in cellular metabolism in response to BNF and light-dark transitions. Particularly, we sought to link changes in protein abundances and pathways to these conditions to dig deeper into the complex relationship between CO_2_ fixation, respiration, and glycogen metabolism that influence temporal separation of photosynthesis and BNF in cyanobacteria.

## Materials and Methods

### Cell growth and experimental conditions

*Crocosphaera sp. subtropica* ATCC 51142 stocks were maintained in ASP2 medium with 17.6 mM NaNO_3_ and 30 µmol photons m^-2^ s^-1^ of continuous light. One-twentieth (5ml) of the *Crocosphaera* 51142 stock was first inoculated to each 250-ml flask containing 100 mL ASP2 medium with or without 17.6mM NaNO_3_ and allowed to grow for 7 days on a shaker at 125 rpm, 30°C, and 30 µmol photons m^-2^ s^-1^ of continuous light. After seven days, cultures were transitioned to alternative 12 h-light/12 h-dark (30 µmol photons m^-2^ s^-1^) diurnal cycles and grown for additional seven days before harvesting. Cells were harvested at 6 h into the light period (L) or 6 h into the dark period (D).

Fifteen (15) ml of cell cultures were collected by centrifugation at 3,220 × g in 15 ml tubes (Corning), washed with 1 ml of 50 mM HEPES-KOH buffer (pH7.5), and then centrifuged again at 10,000 × g at 4°C for 15 min to collect cell pellets. The pellets were resuspended in 200 ul of HEPES/KOH (pH7.8) buffer, supplemented with 1 mM phenylmethylsulfonylfluoride (PMSF) protease inhibitor and homogenized in Precellys VK 0.5 tubes (Bertin Corp., Rockville, MD, USA), 3× at 6000 rpm for 3 x 20 s in each cycle followed by probe sonication. Protein concentration was determined by bicinchoninic acid (BCA) assay (Pierce Chemical Co., Rockford, IL, USA). Following BCA assay, cell lysates corresponding to 200 μg of total protein (equivalent volumes) were ultracentrifuged at 150,000 × g for 20 min to divide proteins into soluble and insoluble fractions. The soluble fractions were acetone precipitated with four volumes of cold acetone and incubated overnight at -20°C, while the insoluble pellets were resuspended in 200μl of HEPES/KOH (pH 7.5), bath sonicated for 5 mins and then acetone precipitated with four volumes of cold (-20°C) acetone. The soluble and insoluble protein pellets were collected by centrifuging at 17,200 × g for 20 min at 4°C, washed 3× with 80% cold (-20°C) acetone and prepared for LC-MS/MS analysis as described below.

### Protein extraction and proteolysis

Both soluble and insoluble pellets were resuspended in 20μl buffer of 8M urea and solubilized by incubating at room temperature with continuous vortexing for an hour, and 0.1% Rapigest (Waters, MA) was added to the insoluble pellets only with the urea solution. The samples were reduced with 10mM dithiothreitol (DTT) at 37°C for 45 min and then cysteines alkylated with iodoethanol mix (195 uL acetonitrile, 4 uL iodoethanol and 1 ul triethyl phosphine) for 45min at 37°C in a dark. After reduction and alkylation, samples were dried in a vacuum centrifuge (Vacufuge Plus, Eppendorf, Enfield, CT) at 45°C, reconstituted in 150 ul of 50 mM ammonium bicarbonate and then digested with trypsin at a 1:25 enzyme to substrate ratio. High pressure digestion was performed using a Barocycler (Pressure Bioscience INC., Easton, MS, USA) at 50°C with 60 cycles, each cycle consisting of 50s at 20,000 PSI and 10s at 1 atm) as described before (22, 23). Digested peptides were cleaned using Pierce Peptide Desalting Spin Columns (Thermo Fisher Scientific, Waltham, MA, USA). Eluted clean peptides were dried in a vacuum centrifuge and reconstituted in 20 μL 0.1% formic acid (FA) in 3% acetonitrile. The peptide concentration was measured using a nanodrop spectrophotometer (ThermoFisher Scientific), adjusted the final concentration in each sample to 1 μg/μl, and 1 μg (1μl) was used for proteomics analysis of the highest concentrated fraction of each sample (either soluble or insoluble), and 0.5ug was injected for the lower concentrated fraction.

### Liquid chromatography tandem mass spectrometry analysis

One μg (1 μl) and 0.5 μg (of the other fraction) of clean peptides were analyzed by reverse-phase HPLC separation using a Dionex UltiMate 3000 RSLC nano system, coupled to a Orbitrap Fusion Lumos mass spectrometer via Nanospray Flex^TM^ electrospray ionization source (Thermo Fisher Scientific) as described previously (23, 24). Briefly, peptides were first loaded into a PepMap C18 trap column (3μm × 75μm ID × 2 cm) (Thermo Fisher Scientific, Waltham, MA, USA), and then separated using a reverse phase 1.7 µm 120 Å IonOptics Aurora Ultimate C18 column (75 µm x, 25cm). The column was maintained at 50 °C, mobile phase solvent A was 0.1% FA in water, solvent B was 0.1% FA in 80% ACN. The loading buffer was 0.1% FA in 2% ACN. Peptides were loaded into the trap column for 5 min at 5 μl/min, then separated with a flow rate of 400 nl/min using a 130 min linear gradient. The concentration of mobile phase B was increased linearly to 8% in five minutes, 27% B in 80 min, and then 45% B at 100 min. After 100 min, it was subsequentially increased to 100% of B at 105 min and held constant for another 7 min before reverting to 2% of B in 112.1 min and maintained at 2% B until the end of the run. The mass spectrometer was operated in positive ion and standard data dependent acquisition (DDA) mode. The spray voltage was set at 2.8 kV, the capillary temperature was 320 °C and the S-lens RF was set at 50. The resolution of Orbitrap mass analyzer was set to 60,000 and 15,000 at 200 *m/z* for MS1 and MS2, respectively, with a maximum injection time of 100 ms for MS1 and 20 ms for MS2. The full scan MS1 spectra were collected in the mass range of 350–1600 *m/z* and the MS2 first fixed mass was 100 *m/z*. The automatic gain control (ACG) target was set to 3 × 10^6^ for MS1 and 1 × 10^5^ for MS2. The fragmentation of precursor ions was accomplished by higher energy C-trap collision dissociation (HCD) at a normalized collision energy setting of 27% and an isolation window of 1.2 *m/z*. The DDA settings were for a minimum intensity threshold of 5 × 10^4^ and a minimum AGC target of 1 × 10^3^. The dynamic exclusion was set at 15 s and accepted charge states were selected from 2 to 7 with 2 as a default charge. The exclude isotope function was activated.

### LC-MS/MS data analysis

LC–MS/MS data were processed with MaxQuant software (Ver 2.0.3.0) (25, 26). Raw spectra were searched against the *Crocosphaera* 51142 protein sequence database obtained from the UniProt (downloaded in July 2022) containing 5403 protein sequences, for protein identification and MS1 based label-free quantitation. The minimum length of the peptides was set at six AA residues in the database search. The following parameters were edited for the searches: precursor mass tolerance was set at 10 ppm, MS/MS mass tolerance was set at 20 ppm, enzyme specificity of trypsin/Lys-C enzyme allowing up to 2 missed cleavages, oxidation of methionine (M) as a variable modification and iodoethanol of cysteine as a fixed modification. The decoy reverse database was considered for data analysis and to control false discovery rate (FDR) and was set at 0.01 (1%) both for peptide spectral match (PSM) and protein identification. The unique plus razor peptides (non-redundant, non-unique peptides assigned to the protein group with most other peptides) were used for peptide quantitation. Only proteins detected with at least one unique peptide and MS/MS ≥ 2 (spectral counts) were considered valid identifications. Label-free quantitation (LFQ) intensity values were used for relative protein abundance comparisons.

### Bioinformatics data analysis

We mainly performed data analysis in Perseus (version 1.6.0.9) (27), Microsoft Excel and data visualized using OriginPro (Version 2022, OriginLab Corporation, Northampton, MA, USA), InteractiVenn (28), Morpheus (https://software.broadinstitute.org/morpheus). Log2-transformed LFQ intensities (protein intensities) were used for further analysis. Coefficients of variations were calculated for raw protein intensities of Hela digest using triplicate runs to determine the reproducibility of LC-MS/MS analysis and label-free quantitation. Data sets were filtered to make sure that identified proteins showed expression in at least three out of four biological replicates of at least one treatment group and the missing values were subsequently replaced by imputation in Perseus that were drawn from a normal distribution. Principal component analysis of treatment effects and biological replicates was performed as described in (29). Multi-sample test (ANOVA) for determining if any of the means of differentiation stages were significantly different from each other was applied to protein data set. For hierarchical clustering and heatmap generation of significant proteins, mean protein abundances of biological replicates were z-scored and clustered using Euclidean as a distance measure for row clustering. Significantly upregulated or downregulated proteins between the treatment groups (± nitrate, L/D, and nitrate × L/D) were determined by ANOVA and a two tallied student’s *t*-test. Differentially expressed proteins were determined with *p*-value <0.05.

### Gene Ontology (GO) analysis

We used three sequence-based function prediction methods: PFP(30), Phylo-PFP(31), and Extended Similarity Group method (ESG) (32) to assign Gene Ontology (GO) terms (33) to protein-coding genes. The PFP algorithm scores GO terms based on Expect (E)-values of sequences with those GO terms retrieved from the UniProt sequence database by PSI-BLAST (34). It then propagates the scores to parental terms on the GO Directed Acyclic Graph (DAG) according to the number of database sequences annotated with parent and child terms. Additionally, based on validation results over a set of benchmark sequences, it assigns a confidence score to GO term predictions. Phylo-PFP, which is an improvement over PFP, improves the performance by incorporating phylogenetic information. The ESG method performs iterative sequence database searches and annotates a query sequence with GO terms. Each annotation is given a probability based on how similar it is to other sequences in the protein similarity graph. To ensure the inclusion of meaningful GO term annotations, we focused exclusively on predictions characterized by high confidence, and all GO terms exceeding confidence score cutoffs of 20,000 for PFP, 0.7 for Phylo-PFP, and 0.7 for ESG were incorporated into our analysis. To improve high-confident prediction, we combined results from three prediction methods and presented the consolidated GO term annotations. Each result file has information about Protein ID, GO ID, Depth, Class, and GO Description. The depth refers to the depth of GO ID in the GO DAG, and class refers to GO functional category (f - molecular function, p-Biological process, c-Cellular Component), and GO Description describes the predicted GO term. The Gene Ontology release 2021-11-16 was used for this analysis.

### Correlation of proteomics and transcriptomics data

Transcriptomics data was obtained from Stockel et al. (35) and matched with our proteomics data. Like the transcriptomics “pooled control”, the proteomics “pooled control” was the average of the intensities of all nitrate-depleted samples. The ratio of D- to the pooled control was then used to calculate the fold change (FC) of proteins. The log_2_(FC) of these values were plotted for both the proteomics and transcriptomics data using scatterplots in OriginPro (Version 2022, OriginLab Corporation, Northampton, MA, USA). A threshold of ±1.5 (or log (foldchange = ±0.58) was used to decide if the fold change was significant enough. The data was divided into various categories from the plots, and colored accordingly, such as those indicated in both proteomics and transcriptomics data with a threshold fold change of ±1.5, or in any one of these, or none. Then, the GO enrichment was performed for these various categories to provide insights into the molecular regulation of several proteins of interest, such as those of the nif gene cluster, proteases, and ribosomal proteins.

## Results and Discussion

### Overview of the proteomic results

The minima and the maxima of O_2_ production occurs around 6-8 hours of the dark and the light periods, respectively (20). Therefore, we choose to examine the proteome at six hours into both the light and dark cycles, to capture the proteome responses during the peak or near-peak of the organism’s metabolic activities related to photosynthesis and nitrogen fixation.

Cells were grown as described previously (36) and as outlined in Figure 1 (see materials and methods for details). Cells were harvested six hours into the light or into the dark cycles and four biological replicates per treatment group were divided into soluble and insoluble preparations by ultracentrifugation before LC-MS/MS analysis. Tryptic peptides were analyzed in a single injection using a 130-min LC method in DDA mode (see Methods for details). MaxQuant search resulted in the identification of 29,946 peptides (Supplementary Table S1) and 2688 proteins (Supplementary Table S2), representing ∼40% of the proteome. Data were filtered to retain a high confidence set of proteins (proteins identified at minimum three of the four replicates in at least one treatment group with LFQ intensity >0 and MS/MS counts >2). This resulted in the identification of 2050 proteins, which were used for downstream bioinformatic and statistical analysis. Of the 2050 proteins, 1321 proteins were shared across all groups and the highest number of proteins were identified under nitrogen-fixing condition in the dark (Figure 2A). Principal component analysis (PCA) showed proteins clustering separately based on treatments with very high concordance between replicates (Figure 2B). In ANOVA comparison, 1370 proteins (*adj. p* value ≤ 0.05) changed due to nitrate, 557 proteins changed due to light and dark, and 361 proteins changed due to the combination of both nitrate and light (Figure 2C, Supplementary Table S3). Hierarchical clustering of significant proteins (ANOVA, *p*-value < 0.05) revealed largest proteome differences in response to nitrate (Supplementary Figure 1A). Nitrogenases were among the most upregulated proteins and the flavoproteins (cce_3835; cce_3833) and heme oxygenase (Ho1; cce_2573), were among the most downregulated proteins in the dark under nitrogen-fixing conditions. The data represented the key proteins in each of the major functional categories that provided important insights about the major metabolic changes during L/D cycles and nitrogen-fixation. The largest number of differentially abundant proteins represented cellular processes such as nitrogen-fixation, photosynthesis and respiration, CO_2_ fixation, carbohydrate metabolism, transport, folding, protein translation, and protein degradation. The predicted proteome of *Crocosphaera* 51142 consists of 5269 open reading frames, of which only 34% are of known function, (8). About 20% of proteins identified in this study hypothetical, uncharacterized or unknown function proteins, highlighting the challenges and need to determine the functions of these hypothetical proteins to better understand the cellular metabolism of *Crocosphaera* 51142 under various growth conditions.

**Figure 1:**
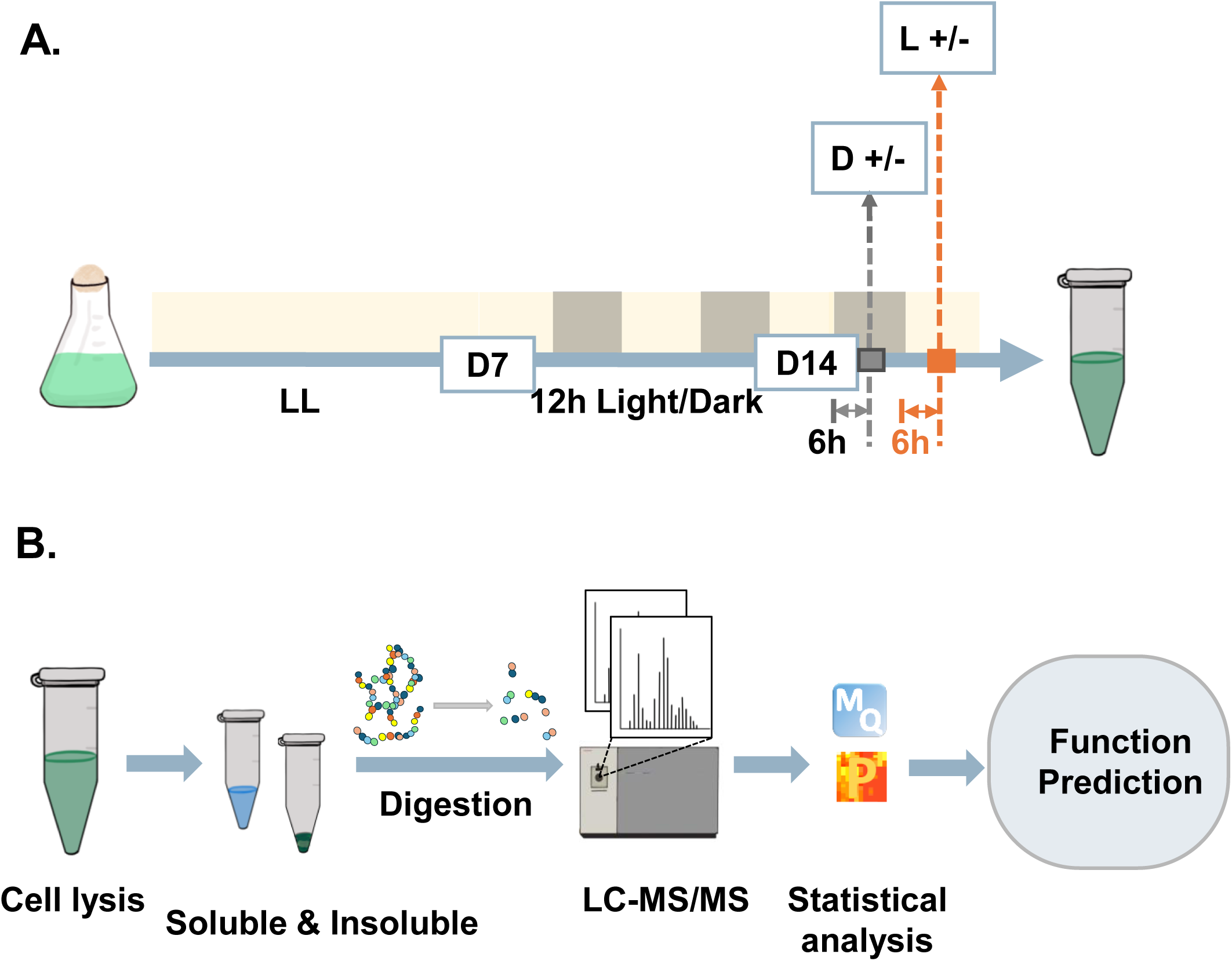
Experimental workflow. (A). Culture growth conditions and sample collection timeline. *Crocosphaera subtropica* ATCC51142 cultures were grown in ASP2 medium without NaNO_3_ (NO ^-^) and with NaNO (NO ^+^) at 30°C under continuous light for seven days. Cultures were then grown for seven days at 12h light/dark (L/D) cycles before harvesting cells at 6 hours into the light (L) and 6 hours into the dark (D). (B). After lysis, cell lysates were separated into soluble and insoluble fractions by differential centrifugation. Soluble lysates were acetone precipitated and insoluble fractions were solubilized with 0.1% Rapigest. Both fractions were digested with trypsin and analyzed by LC-MS/MS with four biological replicates per group. Data processing and statistical analysis were performed using MaxQuant and Perseus respectively. Sequence-based function prediction of the identified proteins was done using PFP, Phylo-PFP and ESG to assign GO terms to the identified proteins.

**Figure 2:**
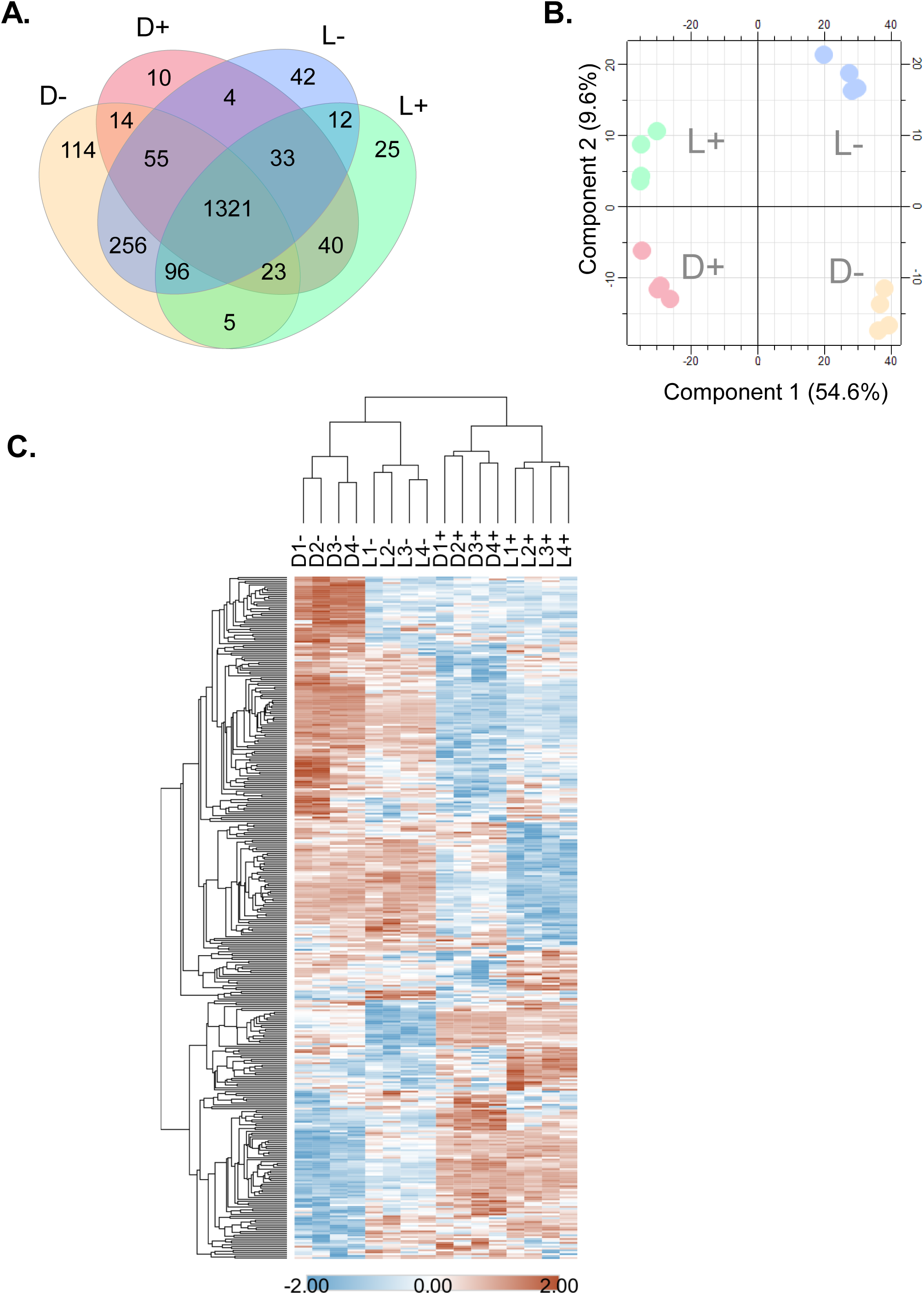
Differentially regulated proteins during diurnal cycles and nitrogen fixation states. (A). Venn Diagram showing protein overlap between D^-^, D^+^, L^-^ and L^+^ conditions. (B). PCA Plot of all the analyzed replicates of D^-^, D^+^, L^-^ and L^+^ samples. The explained variances are indicated in the brackets. (C). Heatmap shows the hierarchical clustering of 361 proteins changing due to interaction of effect of light and dark as well as nitrate. Both rows and columns were clustered by Euclidean distance and average linkage method, column clustering indicates the reproducibility within the replicates for each condition.

To contextualize the differentially regulated proteins across growth conditions, we performed GO enrichment analysis. Figure 3 shows the top enriched GO terms for each treatment condition. Due to the interaction between nitrate and light-dark conditions, nitrogen fixation, molybdenum cofactor binding, tryptophan synthase activity, oxidoreductase activity, and phycobilisome were among the most enriched GO terms. Under the influence of light and dark conditions alone, cellular macromolecule biosynthetic processes, gene expression, primary metabolic processes, and photosynthesis, etc. were enriched. Because of nitrate alone, cellular nitrogen compound biosynthetic process, gene expression, cellular and macromolecular biosynthetic processes were enriched (Supplementary Table S11).

**Figure 3:**
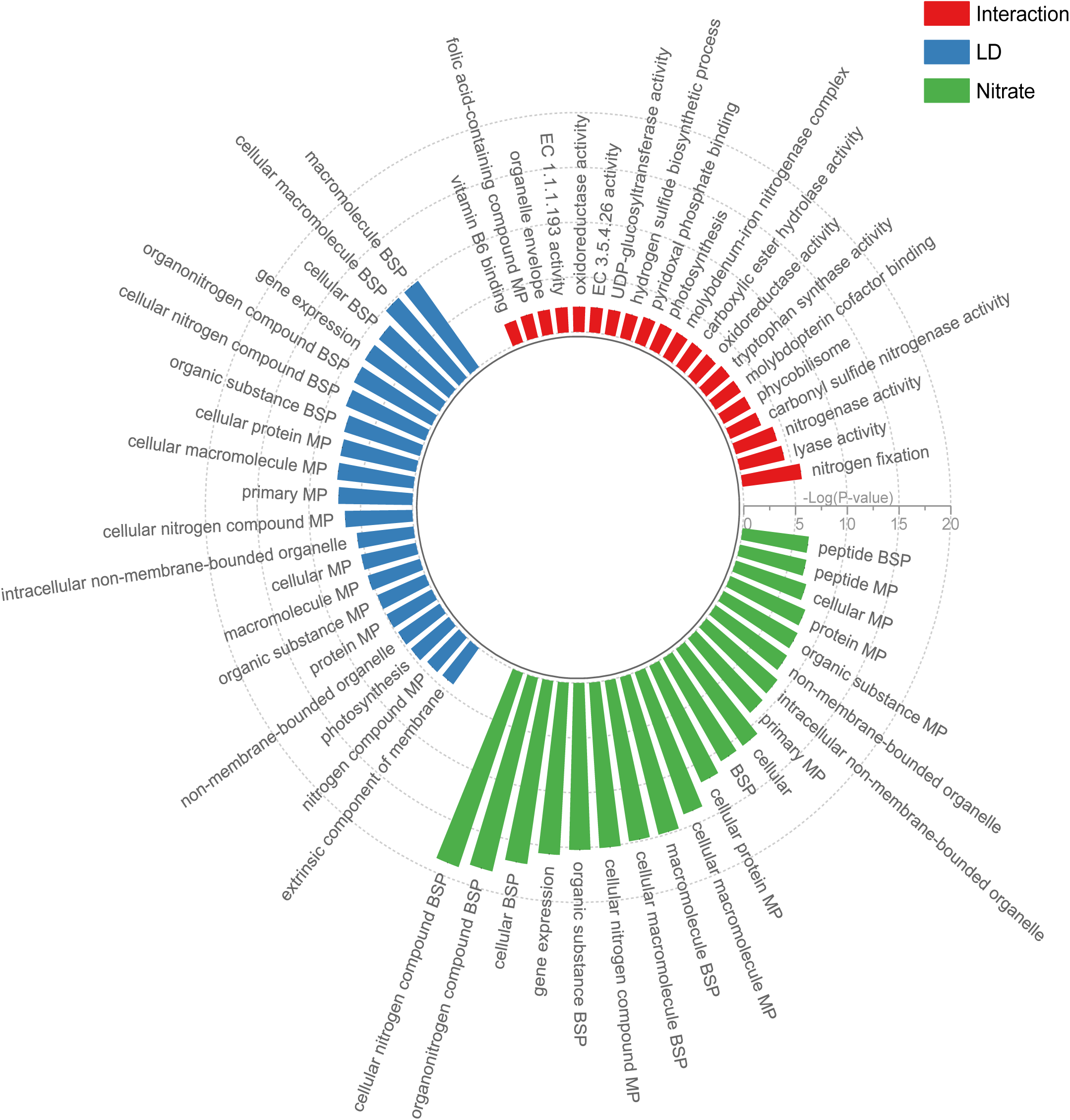
GO classification of significantly changing proteins. Circular bar plot shows the gene ontology classification for the significantly changing proteins due to differences in light and dark (blue), effect of nitrate (green) and interaction of both these conditions (red) (Supplementary Table S11). BSP: Biosynthetic process, MP: Metabolic process, EC 3.5.4.26: diaminohydroxyphosphoribosylaminopyrimidine deaminase, EC 1.1.1.193: 5-amino-6-(5-phosphoribosylamino)uracil reductase.

To determine how the cells grown under nitrogen-fixing growth condition differed from non-fixing growth under L/D cycles, we displayed all quantified proteins as a volcano plots, highlighting the proteins with the highest log transformed fold-change and lowest q-values (Figure 4, A-D). Nitrogenase enzymes together with HupL and PetH were among the top upregulated proteins and CmpA, CmpC, and CmpD were among the top-down regulated proteins in the dark under nitrogen-fixing growth condition compared to non-nitrogen fixing conditions (Figure 4A, Supplementary table S4). Similarly, in the light, PgI, GlpD, MetE and NifH were upregulated in the nitrogen fixing conditions compared to non-nitrogen fixing conditions (Figure 4B, Supplementary Table S5). Comparing the effect of light and dark under nitrogen fixing conditions only, we found that NifK, NifB, Gap were upregulated while Ho1, Ugd and CmpA were downregulated in the dark (D^-^) compared to the light (L^-^) (Figure 4C; Supplementary table S6). On the other hand, under non-nitrogen fixing conditions, psbA4, aphA, kaiC2, TCA Cycle enzymes like talA and glgA1 were upregulated in the dark, compared to acsF, murA and cmpC that were downregulated in the dark compared to light (Figure 4D, Supplementary table S7).

**Figure 4.**
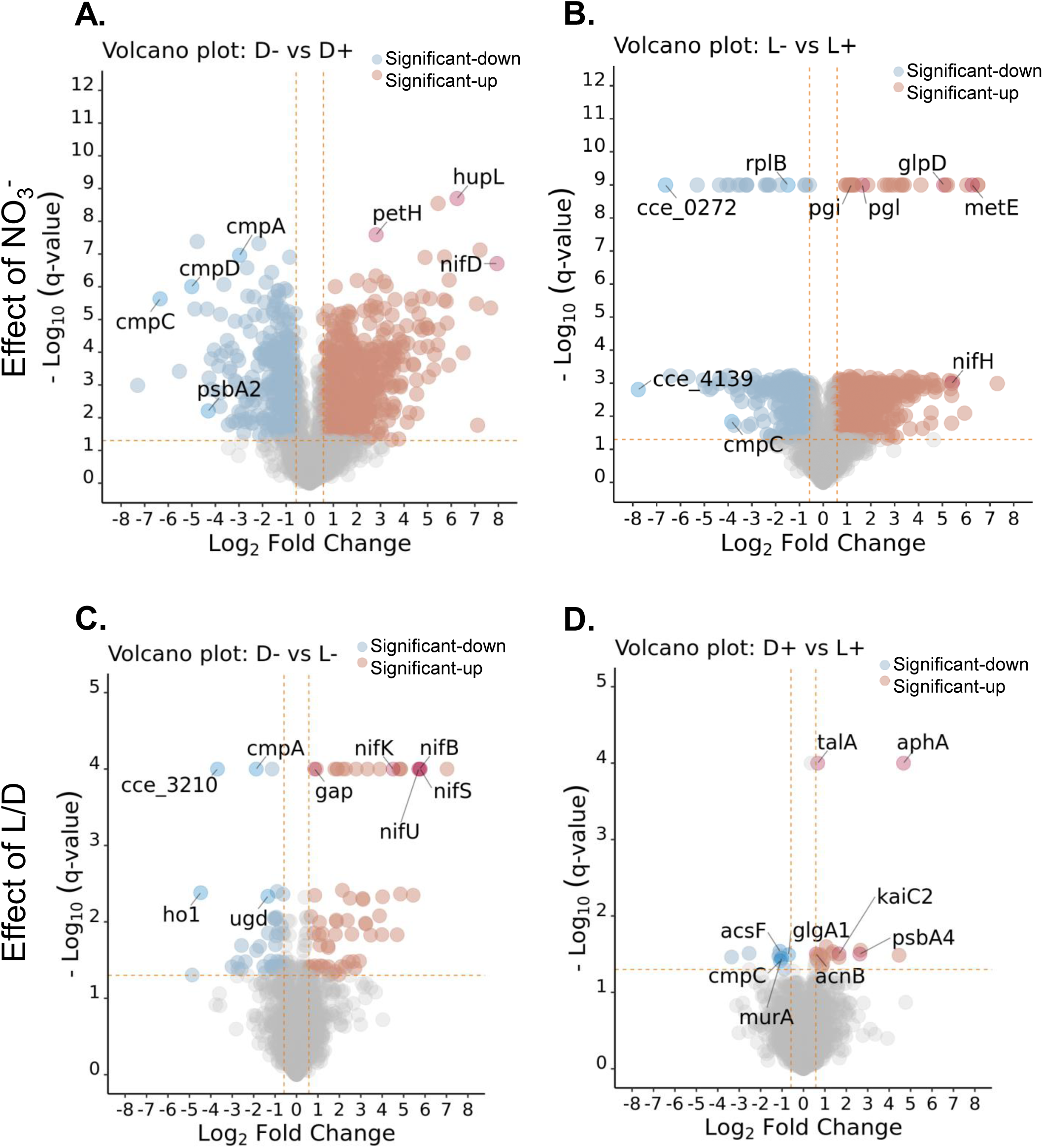
Proteins exhibiting the highest fold change dynamics. Volcano Plots indicate the differential regulation of proteins because of nitrate (A, B) and effect of light and dark (C, D). Significantly changing proteins determined by T-tests with a q-value ≤ 0.05 and threshold log_2_(fold change) =±0.58 are indicated in red (significantly up) and blue (significantly down) in their respective conditions. Proteins exhibiting the highest fold change dynamics in each comparison are specified. Proteins with q=0 were replaced with the lowest possible q-value (or the highest possible –log10(q-value) for the respective dataset.

### Differential regulation of proteins involved in respiration and glycogen metabolism

The cytochrome c oxidase (CoxB1; cce_1977) showed a significant 13-fold higher abundance in the dark under nitrogen-fixing conditions compared to non-fixing conditions (Table 1). In the light cycle, it was 3.8 times. The expression of the nitrogenase enzyme cluster was limited to nitrogen-fixing conditions only (Figure 5A, Table 5B). Nitrogenase expression mirrored with the expression of CoxB1 supporting the potential role of CoxB1 in active respiration to create anoxic condition for nitrogenase activity (15).

**Figure 5:**
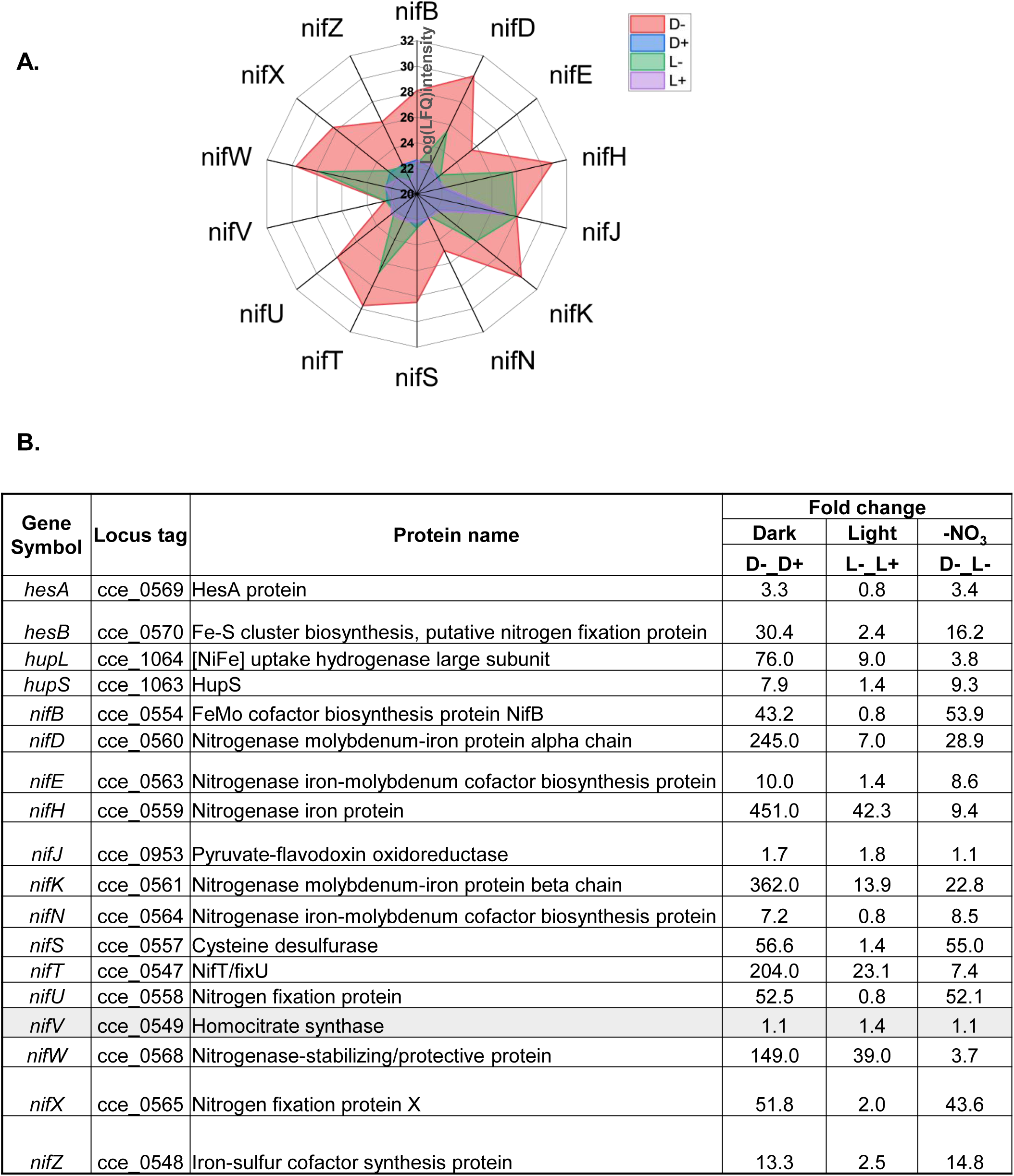
Expression of nitrogenase enzyme clusters. A. Spider chart depicting the differential regulation of enzymes involved in the nitrogenase cluster under various conditions (D^+^/D^-^/L^+^/L^-^). The indicated the log (LFQ intensity) in increasing order. B. List of all the identified nitrogenase clusters and associated genes that play an important role in nitrogen-fixation. All the proteins (except the highlighted ones in grey) are significantly changing in at least one condition.

**Table 1:** Representative list of proteins exhibiting significant changes in at least one pairwise comparison among different experimental conditions (nitrate, light-dark cycles, or interaction between nitrate and light-dark cycles).

Cyanobacteria convert CO_2_ and water into carbohydrates, primarily glucose, during photosynthesis to supporting cellular activities during periods of darkness (37). Enzymes responsible for glycogen synthesis; GlgA1 and GlgA2 were more abundant in the light whereas enzymes for glycogen metabolism; GlgP1 and GlgP2 were more abundant in the dark under nitrogen-fixing condition (Table 1). Active glycogen metabolism generates ATP and creates a low oxygen environment for BNF (6, 9). There are two glycogen debranching enzymes, *glgP*, which is found in the β-cyanobacterial clade (the common ancestor of all cyanobacteria that possessed a β-carboxysome) and *glgX*, which is found in the α-cyanobacteria (which possess α-carboxysome) (7, 8, 38, 39). We identified both enzymes but observed different responses to L/D cycles and nitrate. Among the three GlgP isoforms, GlgP1 (cce_1629) was upregulated whereas GlgP2 (cce_5186) and GlgP3 (cce_1603) were downregulated under nitrogen-fixing condition independent of L/D cycles. The GlgX, on the other hand, was upregulated under nitrogen-fixing condition. These data suggest that glycogen accumulation and degradation is carefully regulated in *Crocosphaera* during BNF.

### Differential regulation of glycolysis, TCA cycle and OPP pathway enzymes

Glycolytic enzymes showed similar abundances between L/D cycles but exhibited stronger response to nitrate (Table 1, Supplementary Figure 2A, Supplementary Table S3). Glucose-6-phosphate dehydrogenase (*Zwf;* cce_2536) directs carbon to OPP pathway and initiates glucose oxidation concomitantly with the generation of NADPH to provide reductants for BNF (40). This enzyme was upregulated in the dark under nitrogen-fixing condition (Table 1). OPP pathway enzymes, including 6-phosphogluconate dehydrogenase (Gnd, cce_3746), OxPPCycle protein (OpcA, cce_2535), transaldolase (TalA, cce_4686), TalC (cce_4208), and 6-phosphogluconolactonase (Pgl, cce_4743) were also upregulated in the dark under nitrogen-fixing condition compared to other growth conditions (Table1). TalA had higher abundance in the dark than in the light but was unaffected by nitrate. TalC was unaffected by the L/D cycle but showed higher abundance without nitrate than with nitrate (Table 1).

TCA cycle enzymes were also in general, more abundant in the dark than in the light, but the effect of nitrate was more pronounced (Table 1, Supplementary Figure 2A). TCA cycle enzyme, isocitrate dehydrogenase (Icd, cce_3202) which converts isocitrate to 2-oxaloacetate (2-OG), which is utilized for the biosynthesis of glutamate via GS-GOGAT cycle (41), was upregulated under nitrogen-fixing growth (Table 1). The succinate semialdehyde dehydrogenase NADP+ (GabD, cce_4228), was lower in nitrogen-fixing cells than non-fixing cells and was also more abundant in the dark than in the light. GabD participates in glutamate metabolism for the conversion of 2-OG to succinate (42). The expression patterns of both Icd and GabD are in agreement with our previous report (16). Increased expression of *icd* transcripts under nitrogen-depleted condition has been reported in other cyanobacteria (41).

Among other TCA cycle enzymes, expression of aconitase (AcnB, cce_3280) was affected by both nitrate as well as L/D cycles independent to each other (Table 1). Under nitrogen-fixing growth, cyanobacteria preferentially utilize alternative nitrogen sources (43). The ammonia (NH_4_^+^) produced during BNF can be trapped by GlnA (cce_4432) and converted to glutamate via GS (GlnA)-GOGAT (GlsF) cycle and incorporated into other amino acids through the activities of aminotransferase or transaminase (44). Increased synthesis of GlnA, GlnB (cce_1775), and GlsA (cce_2244) in the dark cycle under nitrogen-fixation suggest utilization of ammonia produced by BNF for protein synthesis. Indeed, upregulation of carbamoyl-phosphate synthase (CarA, 0902) and CarB (cce_2038) in the dark under nitrogen-fixing conditions suggest the conversion of glutamine into carbamoyl-P, which can be then directed as a substrate for purines and pyrimidine synthesis and for urea cycle. This is supported by the observation that orotate phosphoribosyltransferase (PyrFE, cce_0502) was >2.8 times higher and CTP synthase (PyrG, cce_2923) was >9.3 times higher in the dark under nitrogen-fixing conditions compared to non-fixing conditions (Table 1). ArgB (cce_3224), ArgG (cce_4370), and ArgF (cce_3251), which catalyze reactions to convert carbamoyl-P to citrulline and argininosuccinate through urea cycle were also upregulated in the dark cycle. These results suggest that *Crocosphaera* 51142 not only utilize traditional glycolysis and TCA cycle but also various alternative or modified pathways to optimize its metabolic activities.

### Expression of nitrogenase and hydrogenase enzyme clusters

The acquisition of the nitrogenase complex is a significant evolutionary event that evolved and shaped nitrogen cycling in the ecosystem. *Crocosphaera* 51142 consists of a 35 *nif* genes cluster within a single contiguous region of DNA separated by no more than 3 kb (7). We focused on their expression six hours into the L/D cycles and identified 14 annotated nitrogenase enzymes within the *nif* gene clusters that were differentially regulated and were all exclusively expressed under nitrogen-fixing condition, with actual loss of their identification under non-fixing conditions (Figure 5A, Table 5B). Under nitrogen-fixing growth, while all the nitrogenase enzymes were identified in both the light and the dark cycles, their abundances were significantly higher in the dark than in the light. The level of abundance changes was remarkable with fold changes ranging from 7-fold for NifN to 450-fold for NifH (Table 4B). Such a high level of dynamic range of nitrogenase expression has not been seen in previous proteomic studies. The structural proteins NifHDK were among the most upregulated proteins followed by NifT (cce_0547), and NifW (cce_0568). The 2Fe-2S putative N_2_-fixation related protein (cce_0571) was also one of most highly expressed proteins in the dark with 134-fold higher abundance without nitrate than with nitrate (Figure 5A, Table 5B).

The significant accumulation of NifT and NifW is interesting. While specific function of NifT is unknown (45, 46), this protein is believed to be involved in transporting ammonia produced by nitrogenase within the cell. Previous studies in other strains has revealed that deletion of NifT did not affect diazotrophic growth (47, 48), however nitrogenase activity was higher in the mutant lines than that of the wild type (46), indicating some suppressive effect of NifT on nitrogenase activity. More biochemical studies are needed to determine the functions of NifT in cyanobacteria. NifW is a small nitrogenase stabilizing and protective protein with 116 amino acid residues (13.6 kDa), located within the 35 nitrogenase gene cluster, was 148-fold higher abundance in the dark cycle without nitrate compared to with nitrate. Previous studies have shown higher level of expression of this enzyme towards the end of the dark transition, thus its high abundance six hours into the dark period in the current study is in conformity with its late expression pattern. The FeMo cofactor biosynthesis protein NifB (cce_0554) is a sensitive target for assessing the regulation of BNF (49) and was more than 43-times higher abundance in the dark without nitrate compared to with nitrate. However, its expression was not different in the light cycle between nitrate and non-nitrate growth. The *hesA* (cce_0569) and *hesB* (cce_0570) genes are conserved in diazotrophic cyanobacteria and are within the *nif* gene cluster (46). Proteins encoded by these genes were 3-fold and 30-fold more abundant in the dark under nitrogen-fixing condition compared to N_2_-sufficient conditions (Figure 5A, Table 5B). While their actual function in nitrogenase activity is unknown, recent study indicates their involvement in the efficient production of the MoFe protein (46).

*Crocosphaera* 51142 contains *hupSL* genes for an uptake hydrogenase, and these genes are induced in the dark under nitrogen-fixing growth (15, 50). The small subunit HupS mediates electron transport from the active site of the large subunit (HupL) to redox partners and downstream reactions through a set of Fe-S clusters and the HupL harbors the active site that contains the Ni-Fe (37). The uptake hydrogenase recycle the H_2_ produced as a byproduct during BNF by the nitrogenase activity (51), but it may have other functions such as removing O_2_ from nitrogenase, thereby protecting it from inactivation; and supply reducing equivalents (electrons) to nitrogenase and other enzymes (51, 52). The HupSL enzymes were indeed highly expressed in the dark with HupL showing 76 times higher abundance and HupS showing 8 times higher abundance in the dark under nitrogen-fixing condition than with non-fixing conditions (Figure 5B). These results indicate that the regulation of uptake hydrogenase is influenced by both the absence of light and the absence of nitrate. The gene cluster *kaiABC* encodes circadian clock proteins. Among them, KaiA, KaiC1 and KaiC2 were upregulated under nitrogen-fixing growth compared to non-fixng growth in both the L/D cycles. Noticeably, KaiC2 expression was almost 9-fold higher during the light without nitrate than with nitrate (Supplementary Figure 2B).

### Photosynthesis and CO_2_ fixation

The CO_2_ fixation primarily involves the enzyme Rubisco during the Calvin cycle that results in the synthesis of carbohydrates. As the name goes, Rubisco can also catalyze a reaction with oxygen (O_2_), leading to a process known as photorespiration. Reaction with O_2_ rather than CO_2_ results in loss of some fixed carbon as CO_2_, loss of NH_3_, as well as consumption of ATP, thus decreasing photosynthesis efficiency (53). Catalytic activity of Rubisco is influenced by factors such as the concentration of CO_2_ and the competing oxygenation reaction (54).

Carbonic anhydrase (IcfA1, cce_2257) was upregulated in the light cycle under nitrogen-fixing condition. Bicarbonate transport system substrate binding proteins (CmpA, CmpB, CmpC, and CmpD) were also upregulated in the light cycle, however, unlike IcfA1, they were downregulated in the absence of nitrate (Figure 4C, Table 1). Sodium dependent bicarbonate transporter (SbtA, cce_2939) was downregulated in the dark compared to the light in both nitrogen-fixing as well as non-fixing conditions without nitrate as well as with nitrate. These observations agree with our previous results except SbtA, which was shown to have higher abundance in the dark (16, 55). The Rubisco large subunit (CbbL) was affected by L/D cycles but not by nitrate, however, the small subunit (RbcS) was affected by both the L/D and nitrate. RbcS was down under nitrogen-fixing condition. The expression of glycolate oxidase (cce_3708), on the other hand, was dependent on nitrate not on the L/D cycles. These observations suggest active carboxysome and that activity was influenced by the presence or absence of nitrate.

PSI and PSII proteins were down in the dark compared to the light, and again down without nitrate compared to its presence (Supplementary Table S3). D1 proteins, PsbA1/ PsbA2 were grouped together and showed > 8-fold higher in the dark under nitrogen-fixing conditions than under non-fixing conditions (Table 1). They were >6-fold more abundant in the light under the same nitrate conditions. The effect of L/D cycle on these D1 proteins was minimal. Another PsbA2 protein (cce_3411) and PsbA4 (cce_3477), were higher in the dark compared to the light under both the nitrogen-fixing and non-fixing conditions (Table 1). Higher PsbA4 levels in the dark period have been reported in a previous study (55). The phycobilisome complex was upregulated mostly in the light cycle but was also strongly influenced by nitrate with significant downregulation under nitrogen-fixing conditions. The cytochrome b_6_f complex involved in electron transport chain, linking the PSII and PSI (56) was upregulated in the dark under nitrogen-fixing conditions but was constant between LD cycle under non-fixing conditions.

### Translation, folding, degradation and cellular homeostasis

Multiple metabolic forces including protein synthesis, chaperones, detoxifying proteins, and proteases play crucial roles in maintaining proteome homeostasis in cells (57). Unlike ubiquitin proteasome system in eukaryotes, proteostasis in cyanobacteria is regulated by protein quality control (PQC) system depending on AAA+ proteolytic machines (58). We were interested to determine any changes in proteins involved in these biological processes under L/D cycles and nitrogen. The majority of the identified 30S and 50S ribosomal proteins were downregulated under nitrogen-fixing conditions (Figure 6A, B). Protein homeostasis is tightly controlled by chaperones and proteases (59). The small heat shock protein, HSP20 (HspA3; cce_5270), was 3.5-fold more abundant in the dark and 3.0-fold more abundant in the light when compared between nitrogen-fixing and non-fixing conditions. The chaperone DnaK2 (cce_4004) also showed a significant >7.5-fold higher abundance in the dark and >4.5-fold higher abundance in the light when compared between nitrogen-fixing and non-fixing conditions (Table 1). Nitrogen, not the L/D cycle, had a significant impact on the chaperonin GroEL (cce_1344 and cce_3314). While specific roles of chaperones and heat shock proteins in cyanobacteria have not been thoroughly investigated, increased abundances of HSP20 and GroEL under nitrogen-fixing conditions may suggest cellular stress responses.

**Figure 6.**
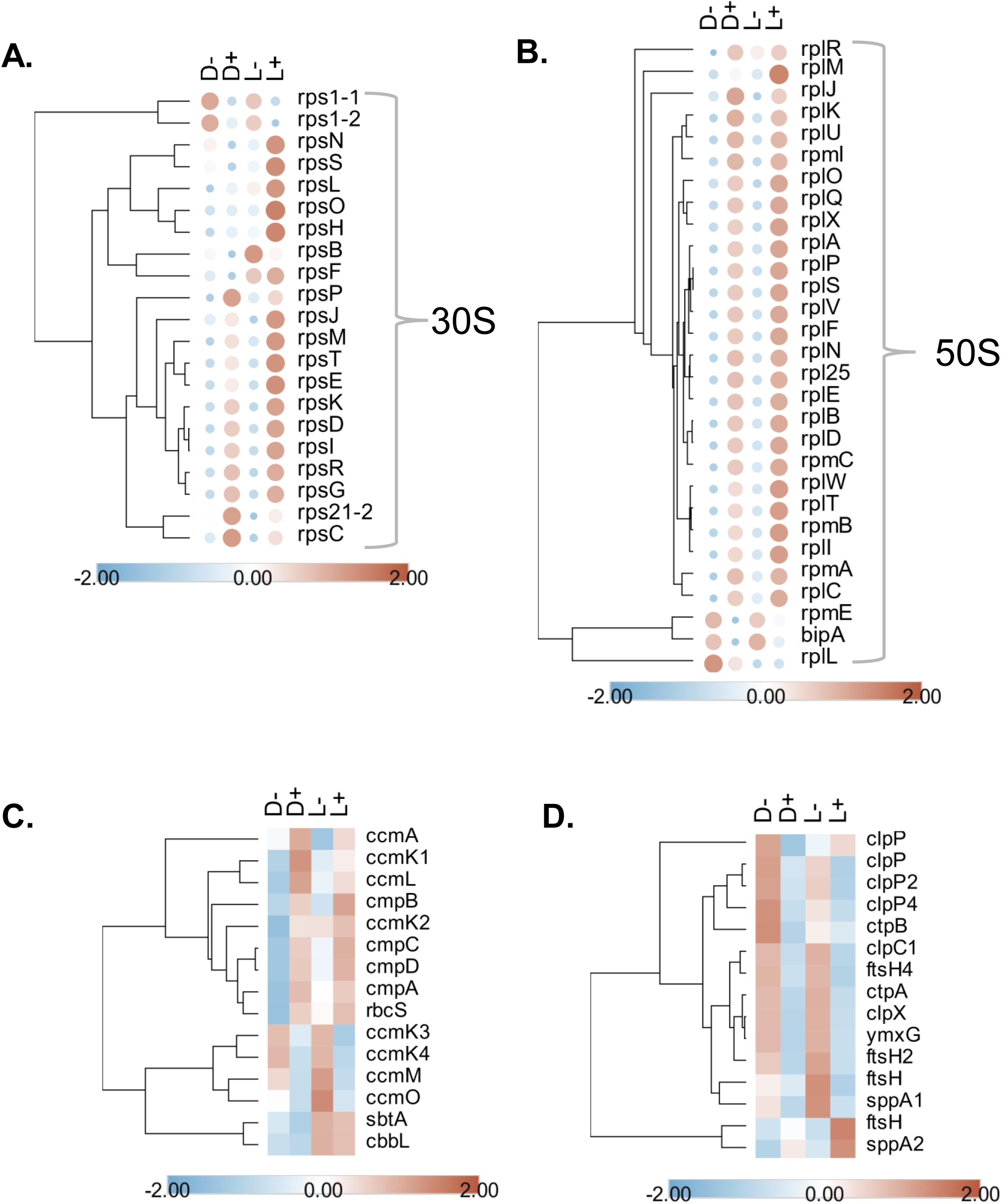
Response of proteins involved in ribosomal subunits, carbon concentrating mechanism and protease degradation. A. Heatmap of proteins involved in 30s (A) and 50s ribosomal subunit (B). Heatmap of proteins involved in carbon concentrating mechanism (C) and proteasomal degradation pathway(D). Hierarchical clustering of the proteins was performed by Euclidean distance measurement and average linkage. The average intensities of all the four replicates for each condition was used to plot these heatmaps.

Protease activities must be tightly regulated in a biological system, particularly in unicellular diazotrophic cyanobacteria, because of their constant shift in metabolic activities between L/D transitions. We identified different classes of proteases including ATP-dependent Clp proteases (ClpX, ClpP, ClpP2, ClpP4, ClpC1), ATP-dependent zinc metalloproteases (Ftsh, FtsH2, FtsH4), CAAX protease, and carboxy-terminal proteases (CtpA, CtpB) (Figure 6D, Table 1). These proteases were, in general, more abundant under nitrogen-fixing conditions than the non-fixing condition in both the L/D cycles. The CtpA (cce_3991), was >5-fold higher under nitrogen-fixing cells compared to non-fixing cells but its expression changed only by 20% between L/D (Table 1). The Clp proteases also showed significant upregulation under nitrogen-fixing condition with ClpP2 showing almost 5-fold higher abundance (Table 1). Among the four zinc metalloproteases, FtsH4 (cce_1593) showed a 2-fold increase in the dark under nitrogen-fixing compared to non-fixing conditions. Our data indicates potential contribution of chaperones, heat shock proteins and diverse classes of proteases in BNF to maintain cellular proteostasis.

### Comparison between transcripts and proteins expressions

It has been demonstrated that in eukaryotic cells, changes in gene transcripts alone can define only one-third of the cell phenotypes, while about 40% can be defined by proteins alone (60), suggesting that both techniques are complementary. There is often a poor correlation between transcripts levels and protein abundances which is attributed to factors such as differences in timing between peak transcript and protein expressions, post-translational modifications, stability, translational efficiency, and protein turnover. Comparing the relationship between mRNA transcript and protein levels is crucial for understanding regulatory processes and functions within cells. While such comparisons have been widely reported in plants (61–63) and animals (64–67), they are less documented in cyanobacteria. Using previously published transcriptomic data (35), which represents transcript level changes at three time points during the dark cycle compared to the light cycle under nitrogen-fixing growth conditions (1h, 5h, and 9h). Our proteomic data represents a single time point at 6 hours into the dark cycle. We identified 868 proteins in our data sets that matched the transcriptomics data. We also identified 158 proteins from those that were significantly changing between D/L cycles under nitrogen-fixing condition that matched the transcriptomics data.

The comparison revealed good correlations between proteomic data and transcriptomic data at one and five hours into the dark cycle, with the strongest correlation observed between one-hour transcript and six-hour protein data (Figure 7, Supplementary Table S8). As expected, there was no correlation between proteomic data and nine-hour transcriptomic data. When comparing significant proteins with corresponding transcripts, a similar correlation was observed under nitrogen-fixing growth conditions. Interestingly, there was a good correlation between transcripts of nitrogen fixation (*nif)* genes and proteins at all time points, suggesting that *nif* transcripts are stable and present throughout the dark cycle (Figure 7).

**Figure 7:**
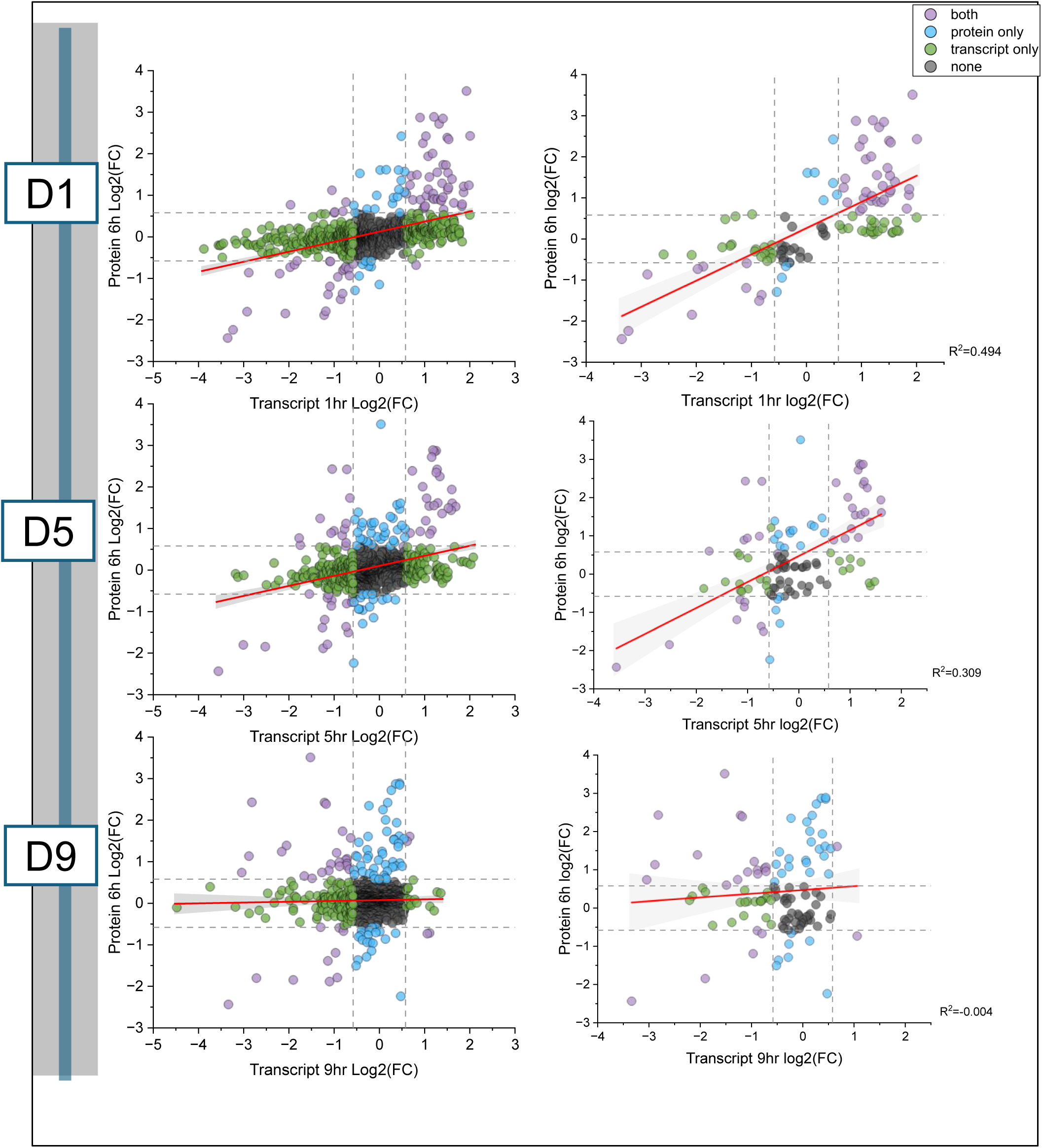
Comparison of proteomics and transcriptomics data. Scatterplots show the comparison of proteomics and transcriptomics data. The left panel shows the correlation of all the identified proteins in D-condition that were common between the proteomic and transcriptomic data, and the right panel shows the significantly changing proteins in D-compared to the pooled control. Proteomics data at 6hours into the dark was compared with transcriptomics data 1 hour, 5 hours and 9 hours into the dark respectively. The color of the data points indicates if the proteins were identified with a threshold fold change of ±1.5 (or log (foldchange= ±0.58) under both conditions (purple), protein only(blue), transcript only(green) or none(grey).

To focus on the nitrogenase enzymes, we separately plotted them with *nif* transcripts (Supplementary Figure 3). The proteomic data showed good correlation with the five hours transcriptomic data. However, as expected one and nine hours of *nif* transcripts showed poor to no correlation. The comparison, though highlights timing differences between the peak of transcripts and proteins in *Crocosphaera* 51142, many transcripts peak at early dark phases and remain stable several hours into the phase, suggesting that the relationship between transcripts and proteins depends on individual or groups of transcripts and proteins involved in specific pathways. This comparison sheds light on the regulation of gene expression and protein abundance to coordinate cellular processes, particularly under nitrogen-fixing growth conditions.

### Conclusions

Our analysis provides a global view of how nitrogen and light-dark phases impact the *Crocosphaera* 51142 proteome. Results demonstrate significant modulation of metabolic pathways by cells to adapt to transitions between light-dark phases under nitrogen-fixing or non-fixing conditions. Regulation of CoxB1 and GlgP1 in synchronization with nitrogenase and uptake hydrogenase suggests that cells maintained suboxic condition for nitrogenase activities by higher levels of respiration and glycogen metabolism. Previous transcriptomics and proteomics studies in *Crocosphaera* 51142 and other strains have also reported significant upregulation of *nif* genes and proteins ranging from 10-40-fold during the dark cycle under nitrogen-fixing conditions. However, our current data reveals a dynamic range of expression levels for Nif proteins, ranging from a 7-fold to a dramatic 450-fold increase, specifically after six hours into the dark cycle under nitrogen-fixing conditions, is remarkable, and has not been observed before. Such a remarkable and previously unreported dynamic range of nitrogenase expression in response to nitrogen-fixing conditions during the dark cycle is important and may contribute to the understanding of the regulatory mechanisms governing BNF in cyanobacteria and may have broader implications for the optimization of nitrogen-fixing cyanobacteria in various applications, including biotechnology and environmental management.

## Data Availability

All the raw LC-MS/MS data are deposited in MassIVE data repository (massive.ucsd.edu) with MASSIVE-ID: MSV000094471 All the mapped spectra for the annotated peptides in the global dataset are available in the MS Viewer repository and can be accessed using the search key **q5zmiw9ebd** and by using the following URL: https://msviewer.ucsf.edu/prospector/cgi-bin/mssearch.cgi?report_title=MS-Viewer&search_key=q5zmiw9ebd&search_name=msviewer

## Acknowledgements

All LC-MS/MS experiments were performed at the Purdue Proteomics Facility in the Bindley Bioscience Center of Purdue University. We thank Rodrigo Mohallem and other members of the Aryal lab for discussion and feedback in data analysis and interpretation.

## Authors contributions

**Punyatoya Panda**: Culture growth, treatments, proteomics sample preparation, LC-MS/MS data acquisition, data analysis, writing original draft, review, and editing. **Swagarika J. Giri**: GO annotation, data analysis, writing original draft, review, and editing. **Louis Sherman**: conceptualization, project supervision, data interpretation, review, and editing. **Daisuke Kihara**: conceptualization, data analysis, interpretation, manuscript editing, fund acquisition. **Uma K. Aryal**: conceptualization, methodology, data collection, data analysis and interpretation, writing original manuscript, review, and editing, fund acquisition, project supervision.

## Notes

The authors declare no competing financial interests.

## Abbreviations

LC-MS/MS: Liquid Chromatography-Tandem Mass Spectrometry
RSLC: Rapid Separation Liquid Chromatography
HEPES-KOH: 4-(2-hydroxyethyl)piperazine-1-ethanesulfonic acid potassium salt
ATCC: American Type Culture Collection
L^-^: Cells harvested six hours into the light cycle under nitrogen fixing condition.
L^+^: Cells harvested six hours into the light cycle under nitrogen non-fixing condition.
D^-^: Cells harvested six hours into the dark cycle under nitrogen fixing condition.
D^+^: Cells harvested six hours into the dark cycle under nitrogen non-fixing condition.
BNF: Biological Nitrogen Fixation
PSI and PSII: Photosystem I and Photosystem II
GO: Gene Ontology
CO_2_: Carbon dioxide
N_2_: Dinitrogen
Nif: Nitrogenase enzymes
O_2_: Oxygen
PQC: Protein Quality Control
ATP: Adenosine Triphosphate
AAA+ Protease: ATP Associated Protease
NADPH: Nicotinamide Adenine Dinucleotide Phosphate

