## Supplementary figures and images for "Proteomic changes orchestrate metabolic acclimation of a unicellular diazotrophic cyanobacterium during light-dark cycle and nitrogen fixation states"

### Supplementary Figures

Supplementary Figure 1

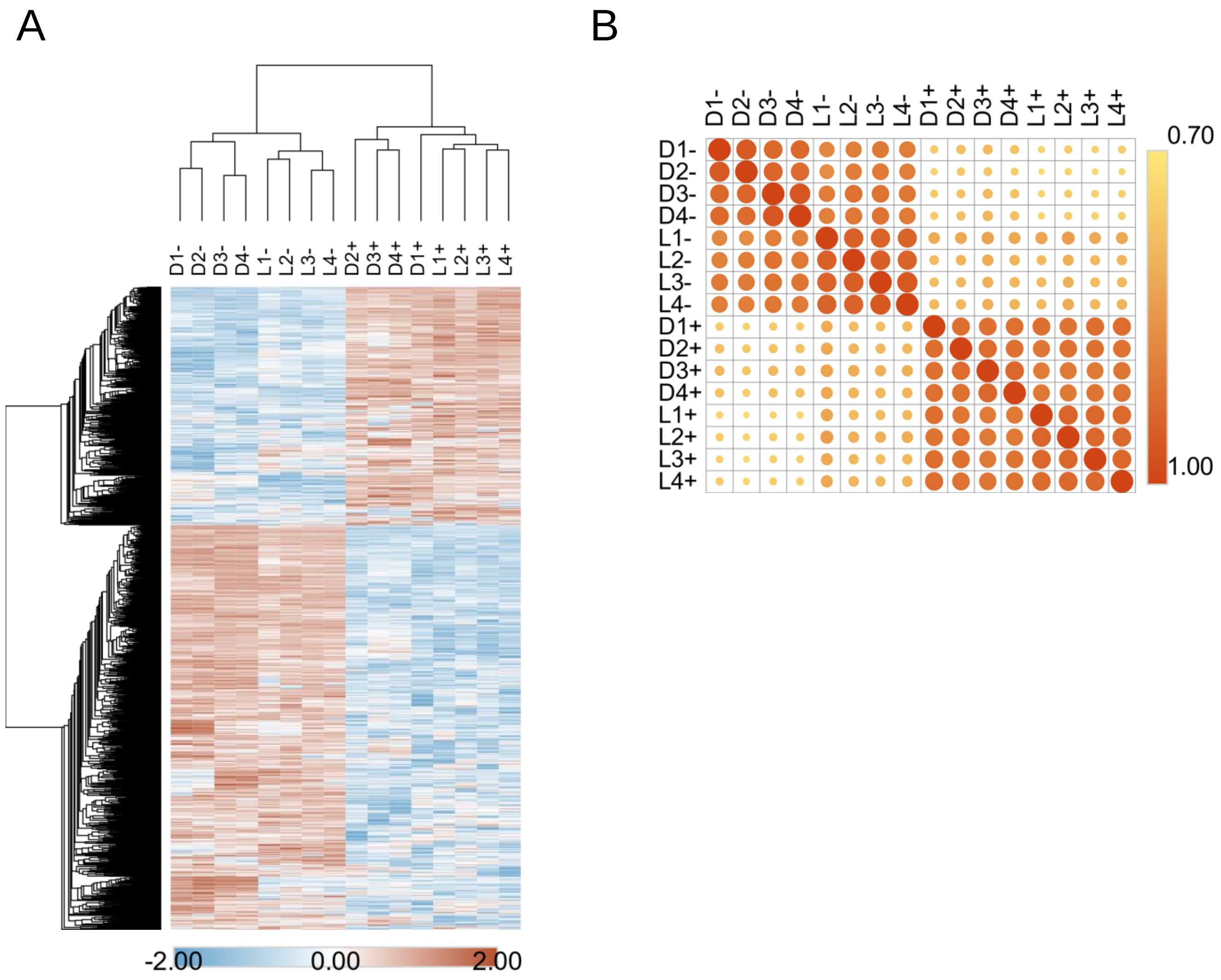

#### Supplementary Figure 2

A.

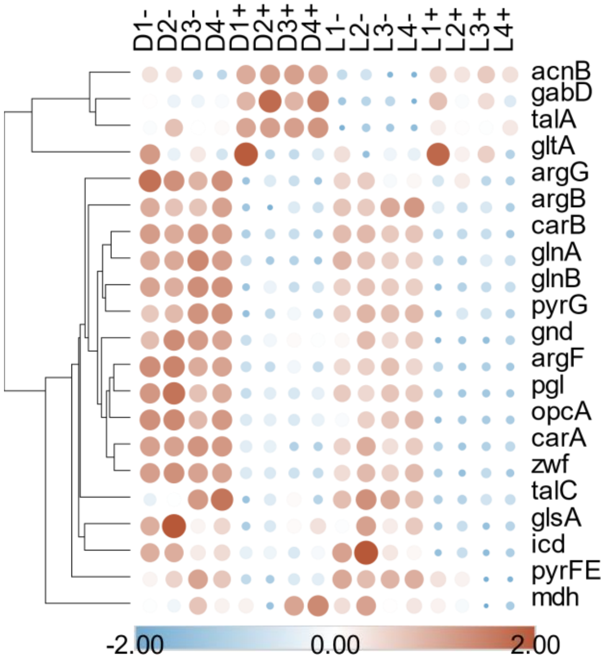


B.

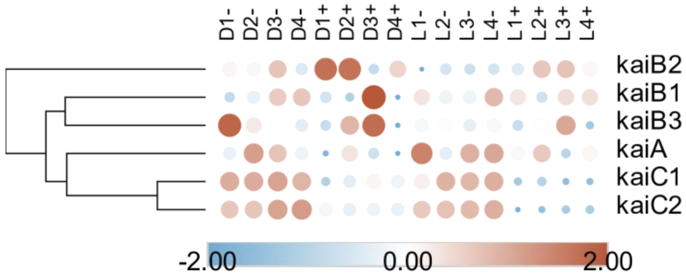

#### Supplementary Figure 3

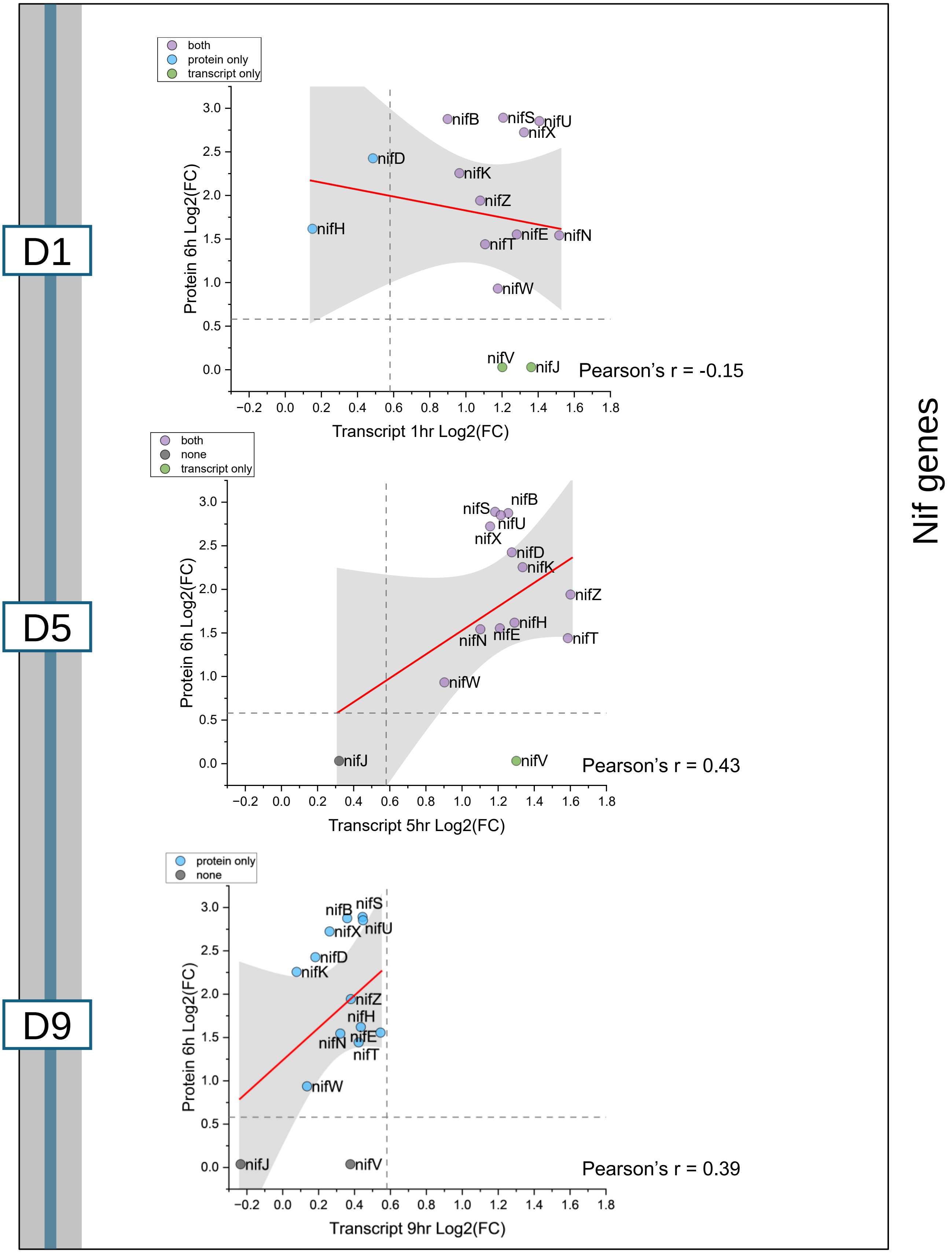
